# Deep-learning-enabled multi-omics analyses for prediction of future metastasis in cancer

**DOI:** 10.1101/2025.05.16.654579

**Authors:** Xiaoying Wang, Maoteng Duan, Po-Lan Su, Jianying Li, Jordan Krull, Jiacheng Jin, Hu Chen, Yuhan Sun, Weidong Wu, Kai He, Richard L. Carpenter, Chi Zhang, Sha Cao, Dong Xu, Guangyu Wang, Lang Li, Gang Xin, David P. Carbone, Zihai Li, Qin Ma

## Abstract

Metastasis remains the leading cause of cancer-related mortality, yet predicting future metastasis is a major clinical challenge due to the lack of validated biomarkers and effective assessment methods. Here, we present EmitGCL, a deep-learning framework that accurately predicts future metastasis and its corresponding biomarkers. Based on a comprehensive benchmarking comparison, EmitGCL outperformed other computational tools across six cancer types from seven cohorts of patients with superior sensitivity and specificity. It captured occult metastatic cells in a patient with a lymph node-negative breast cancer, who was declared to have no evidence of disease by conventional imaging methods but was later confirmed to have a metastatic disease. Notably, EmitGCL identified *HSP90AA1* and *HSP90AB1* as predictable biomarkers for future breast cancer metastasis, which was validated across five independent cohorts of patients (n=420). Furthermore, we demonstrated YY1 transcription factor as a key driver of breast cancer metastasis which was validated through *in-silico* and CRISPR-based migration assays, suggesting that YY1 is a potential therapeutic target for deterring metastasis.

## Main

Advances in multi-modality therapy such as surgery, radiation therapy, chemotherapy, targeted therapy and immuno-therapy have improved survival outcomes in cancer patients, but metastatic disease remains a major challenge^1,2^. Clinical studies have primarily focused on identifying predictive biomarkers for metastatic risk such as (*i*) imaging-based markers, including radiomic features extracted from MRI and PET scans, (*ii*) HER2 enrichment^3^ which has been identified as a predictor of brain metastasis in breast cancer patients, and (*iii*) circulating tumor cells, which emerging studies suggest may serve as predictive markers for distant organ involvement^4^. However, most biomarkers were identified based on detectable metastatic cells^5-7^, and they fail to capture cancer cells in metastatic sites that have already spread but remain undetectable due to their rarity^8^ (i.e., **occult metastatic cells**) and those with the potential to initiate metastasis at the primary site^9^ (i.e., **metastatic precursor cells**). As a result, existing biomarkers are often prone to overdiagnosis (i.e., false positive) or to missing actual metastases (i.e., false negative), which can lead to unnecessary treatments or delayed diagnosis^10^. Computational studies identify predictive biomarkers through differentially expressed genes based on bulk RNA-seq data^11-14^. Still, no common biomarkers across different cohort groups of patients have been identified, limiting their generalizability and effectiveness^15,16^. These underscore the need for more sensitive and specific biomarkers to determine whether a cancer patient with no detectable metastasis is likely to develop it over time (i.e., **future metastasis**)^6^.

With advancements in single-cell RNA sequencing (**scRNA-seq**), researchers can now detect various cell types and states, including rare subpopulations, and characterize cellular heterogeneity of cancer at high resolution^17,18^. However, identifying metastatic precursor cells with high metastatic potential within primary tumors and occult metastatic cells at metastatic sites remains challenging for existing computational methods and pathological examination techniques^19^. This limitation arises from (*i*) low sensitivity combined with high FPRs in identifying occult metastatic cells and (*ii*) the inability to trace the dynamic metastatic transition between precursor and occult metastatic cells.

Advancements in artificial intelligence (AI) offer a potential solution by enabling deep feature extraction and pattern recognition in single-cell omics data analysis^20,21 22^. In this study, we developed **EmitGCL** (Futur<u>e</u> <u>m</u>etastasis prediction and <u>i</u>nnovation of <u>t</u>herapeutics based on <u>g</u>raph <u>c</u>ontrastive <u>l</u>earning), which is a knowledge-aware contrastive learning framework designed to predict future metastasis and corresponding biomarkers. We demonstrated that EmitGCL achieves the highest true positive rate (TPR) in detecting occult metastatic cells from metastatic patients based on scRNA-seq data. It outperformed the second-best tool by improving the TPR by 12.44% using 15 scRNA-seq data across pancreas, nasopharyngeal, papillary thyroid, and head and neck cancers. In a very representative breast cancer case, EmitGCL effectively captured occult metastatic cells in the lymph node-negative breast cancer patient, which conventional imaging methods failed to detect but were later confirmed as metastatic. Meanwhile, it did not detect occult metastatic cells in two non-metastatic lung cancer patients, unlike other computational tools, thereby maintaining a 0% false positive rate (FPR) and minimizing the misclassification of non-metastatic patients as metastatic. This enables more precise risk stratification and facilitates timely therapeutic interventions.

To demonstrate the clinical impact of EmitGCL, we identified *HSP90AA1* and *HSP90AB1*, through the attention mechanism in the EmitGCL framework, as markers of future metastasis in breast cancer. These genes exhibited 11%–16% higher accuracy compared to approaches relying on differentially expressed gene analysis in the identification of clinically evident metastatic cells. Two biomarkers were validated using RNA-seq data from 420 breast cancer patients across five cohorts in the UCSC Xena^**23**^ database, demonstrating them as novel, detectable biomarkers for metastasis prevention. In addition, we identified YY1 as a therapeutic target for mitigating metastasis in breast cancer by analyzing ChIP-seq data to map its binding sites to the promoter regions of biomarker genes. The role of YY1 was further validated through an *in-silico* transcription factor (TF) knockout using CellOracle^24^, CRISPR-based perturbation, migration assays, and four independent clinical cohorts. These findings support YY1 as a viable therapeutic target in clinical applications. Biomarkers and TF demonstrate the immediate clinical impact of EmitGCL in advancing outcome improvement and therapeutic development.

## Results

### Dataset characteristics and system overview

EmitGCL is a knowledge-aware contrastive learning approach to identify metastatic precursor cells and biomarkers. The model emphasizes AI modeling, computational analysis, and experimental and clinical validation to ensure the accuracy and applicability of its findings **(Fig. 1a and Supplementary Fig.1)**. To improve the signal-to-noise ratio in the matched scRNA-seq from primary and metastatic sites, we constructed a heterogeneous graph where nodes represent cells and genes, and unweighted edges indicate the presence of genes in cells **(Fig. 1b)**. Based on the hypothesis that metastatic precursor cells are more similar to occult metastatic cells within the same clonotype than to other cells, we predict future metastasis more reliably than existing metastatic risk prediction methods by: (*i*) enhancing TPR, which indicates more accurately identifying occult metastatic cells from metastatic patients. This is achieved through a graph contrastive learning approach, which distinguishes subtle differences in the same tumor clonotype between primary and metastatic sites **(Fig. 1c)**, (*ii*) reducing FPR, which indicates minimizing misclassification of non-metastatic patients as metastasis. This is accomplished by integrating prior knowledge from metastasis-related biological process databases **(Fig. 1d)**, and (*iii*) identifying metastatic precursor cells by tracing occult metastatic cells through site-specific contrastive learning **(Fig. 1e)**.

**Fig. 1.**
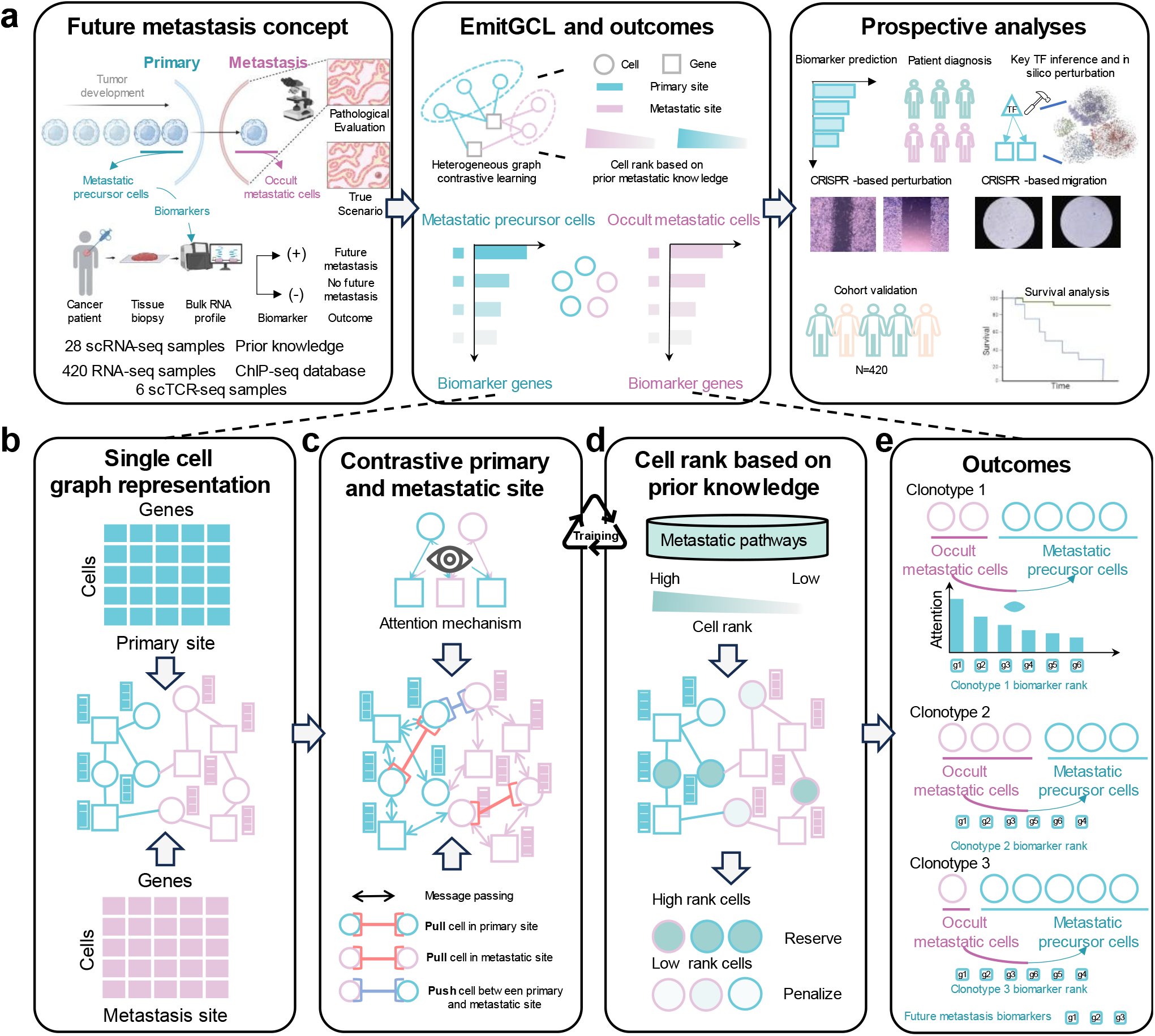
Workflow of the EmitGCL framework. **a**, Various omics datasets were collected from public databases and literature. EmitGCL was trained using heterogeneous graphs with knowledge-aware contrastive learning to predict precursor metastatic cells and biomarkers. Multiple genes and TFs were identified as markers or therapeutic targets for future metastasis prediction, which were validated using experimental and clinical cohorts. Blue means primary site. Pink means metastatic sites. Circles represent cells, and squares represent genes. **b**, scRNA-seq data from both were collected to construct a heterogeneous graph. **c**, The contrastive learning process amplifies differences between primary and metastatic sites within the same clonotype. **d**, The metastatic pathways were involved in model training by penalizing cells with low metastatic potential scores. **e**, The system outputs include metastatic precursor cells and biomarkers in the primary tumor, as well as occult metastatic cells at metastatic sites.

We collected all publicly available matched scRNA-seq datasets from primary and metastatic sites, resulting in 33 paired datasets across breast, pancreas, nasopharyngeal, papillary thyroid, and head and neck cancers. These datasets were derived from seven published studies, involving 28 patients and a total of 516,093 cells. Of these, 215,443 cells are from primary sites, and the other 300,650 are from metastatic sites. To predict future metastasis, it is crucial to identify metastatic precursor cells as they will serve as the metastatic niche of future metastasis development. However, a clinically applicable biomarker that accurately predicts precursor cells requires precise identification of occult metastatic cells, as they provide the key linkage between precursor cells and actual metastatic progression. Furthermore, identifying biomarkers of metastatic precursor cells provides clinically applicable detection methods for future metastasis assessment. To systematically evaluate the performance of occult metastatic cell identification, we first applied our model, which integrates metastatic prior knowledge, to identify occult metastatic cells. Based on the model’s predictions, we then constructed a benchmarking framework to assess performance in terms of TPR and FPR. Additionally, to validate the clinical impact of our model’s inferred biomarkers in metastatic precursor cells, we collected data from 420 breast cancer patients with metastasis across five cohorts **(Fig. 1a)**. To validate TF candidates that drive metastasis, we integrated CHIP-seq data and performed *in-silico* TF knockout experiments using CellOracle, CRISPR-based perturbation, migration assays, and survival analysis across four independent clinical cohorts.

### Evaluation of computational methods for identifying occult metastasis cells

Firstly, we assessed the performance of EmitGCL in identifying occult metastasis cells using matched scRNA-seq data from primary and metastatic sites across six cancer types **(Fig. 2a)**. Among the 28 patients with 33 paired scRNA-seq datasets, metastasis was observed in five of the six cancer types (**Supplementary Data 1**). The exception was a case of lung cancer from a single patient with two lymph nodes, which did not exhibit metastatic progression. Pancreatic cancer was associated with liver metastasis, while the other four types (breast, nasopharyngeal, papillary thyroid, and head and neck) demonstrated lymph node metastasis **(Fig. 2b and 2c)**. To validate our model, we compared EmitGCL against four published methods based on their performance reported in the MarsGT^25^ study. MarsGT demonstrated the best performance for rare cell identification in that study. Seurat^26^ serves as the baseline for cell clustering. GiniClust^17^, was shown as the second-best tool for rare cell identification using clustering methods, while CellSIUS^27^ was the second-best tool for rare cell identification using classification methods.

**Fig. 2.**
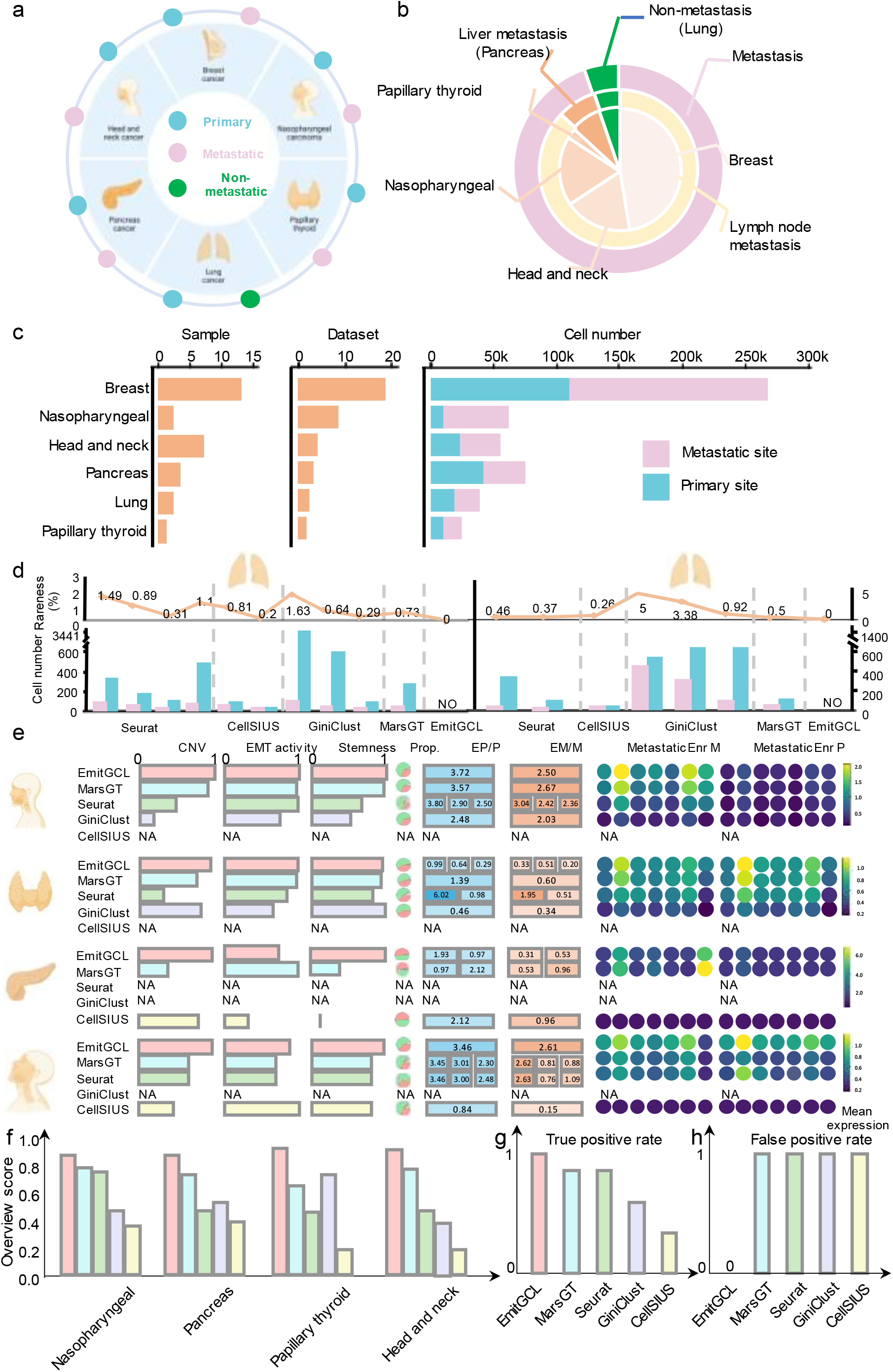
Data collection and computational evaluation of the model for occult metastatic cell prediction. **a**, Schematic depicting the study design (created with BioRender.com). Published paired scRNA-seq data from six cancer types were used. Blue circles represent primary sites, pink circles represent metastatic sites, and green circles represent cancer non-metastasis. **b**, Pie chart showing data distribution. The outer circle represents metastasis versus non-metastasis, the middle segment shows metastatic sites, and the inner circle depicts the distribution of cancer types. **c**, Bar plots summarizing the number of samples, datasets, and cells collected, categorized by primary organ and metastatic sites. **d**, False positive evaluation of occult metastatic cell identification in lung cancer datasets without metastasis. The bar plots represent the number of identified metastatic precursor cells in primary tumors and associated metastatic cells in the lymph node. The line plots display the proportion of metastatic cells among all cells in the metastatic site. **e**, Ture positive analysis of metastatic cell identification across four cancer types. Metrics include CNV, EMT activity, stemness, cell group proportion, and enrichment scores of metastasis-related pathways within metastatic precursor and occult metastatic cells. These evaluations are compared across different methods for each cancer type. **f**, Overall performance score for metastatic cell identification in each cancer type. The y-axis indicates the proportion of correctly identified metastatic datasets relative to the total datasets for each cancer type. **g**, Accuracy evaluation of metastatic cell identification across all datasets. Bar plots represent accuracy scores for each method. **h**, False positive evaluation of metastatic cell identification across all datasets. Bar plots depict FPR for each method.

To evaluate the FPR in identifying occult metastatic cells, we used two paired scRNA-seq datasets from primary lung cancer and lymph node sites that did not show metastasis. The results indicated that although all other tools misclassified non-occult metastatic cells as occult metastatic cells in these datasets, EmitGCL did not identify any occult metastatic cells, demonstrating its low FPR in identifying occult metastatic cells **(Fig. 2d, Supplementary Data 2)**. Furthermore, to validate the TPR of occult metastasis identification, we evaluated EmitGCL alongside other tools across pancreas, nasopharyngeal, papillary thyroid, and head and neck cancers that have metastasized. We visualized several key biological features in the identified occult metastatic cells, including copy number variation (CNV)^28,29^, epithelial-to-mesenchymal transition (EMT) activity scores^30^, tumor stemness scores^30^, and metastasis-related pathway enrichment scores^31^. EmitGCL consistently outperformed other tools, with the identified occult metastatic cells showing higher performance across these metrics **(Fig. 2e, Supplementary Fig. 2, and Supplementary Data 3-8)**. Next, we calculated the rate of occult metastatic cell detection across all datasets for the above four cancer types, which demonstrated that EmitGCL has a higher TPR in identifying occult metastatic cells compared to other methods **(Fig. 2f)**. To provide an overall summary of performance, we compared the TPR and FPR across all datasets **(Methods)**, showing that it effectively reduces FPR while maintaining high TPR **(Fig. 2g, Fig. 2h, and Supplementary Data 9)**.

To further assess EmitGCL’s robustness, we ran the model five times for each cancer type and evaluated consistency using the adjusted rand index (**ARI**), where a higher ARI indicates greater model robustness. It achieved ARI scores higher than 0.95 across all runs, confirming its robustness and reproducibility **(Supplementary Fig. 3)**. Additionally, we validated that EmitGCL-identified biomarker genes, including known metastasis-related genes like *B2M*^32^and *MALAT1*^33^, were consistently detected across primary and metastatic sites, aligning with previous finding **(Supplementary Fig. 4 and Supplementary Data 10)**.

### Validation of occult metastatic cell prediction based on clinical trace data

In the benchmark section **(Fig. 1e and Supplementary Fig. 2)**, we demonstrated that EmitGCL can fully identify patients with metastasis across four cancer types. To further validate our model’s ability to detect metastasis, we analyzed a representative case: paired scRNA-seq data from a breast cancer patient with a primary tumor and two lymph nodes (LN-1 and LN-2), which were initially deemed negative but later confirmed as metastatic, despite being undetectable by conventional pathological evaluation^34^. EmitGCL identified cell cluster 2 in LN-1 and cell cluster 4 in LN-2 as metastatic precursor and occult metastatic cells **(Fig. 3a)**. The corresponding cell counts in the primary and metastatic sites were 995 and 158 in LN-1, and 1,109 and 23 in LN-2, respectively **(Fig. 3b)**. The proportion of occult metastatic cells in the metastatic sites was less than 3%, highlighting the rarity of these ‘ population **(Fig. 3c)**. Further analysis revealed that these clusters exhibited high CNV scores and cancer cell stemness scores, distinguishing them from other cell clusters in LN-1 and LN-2 **(Fig. 3d and 3e)**. Additionally, the EMT scores in the identified occult metastatic cells were significantly higher than those in the metastatic precursor cells, providing further evidence that cell cluster 2 in LN-1 and cell cluster 4 in LN-2 represent true occult metastatic populations **(Fig. 3f and Supplementary Data 11)**.

**Fig. 3.**
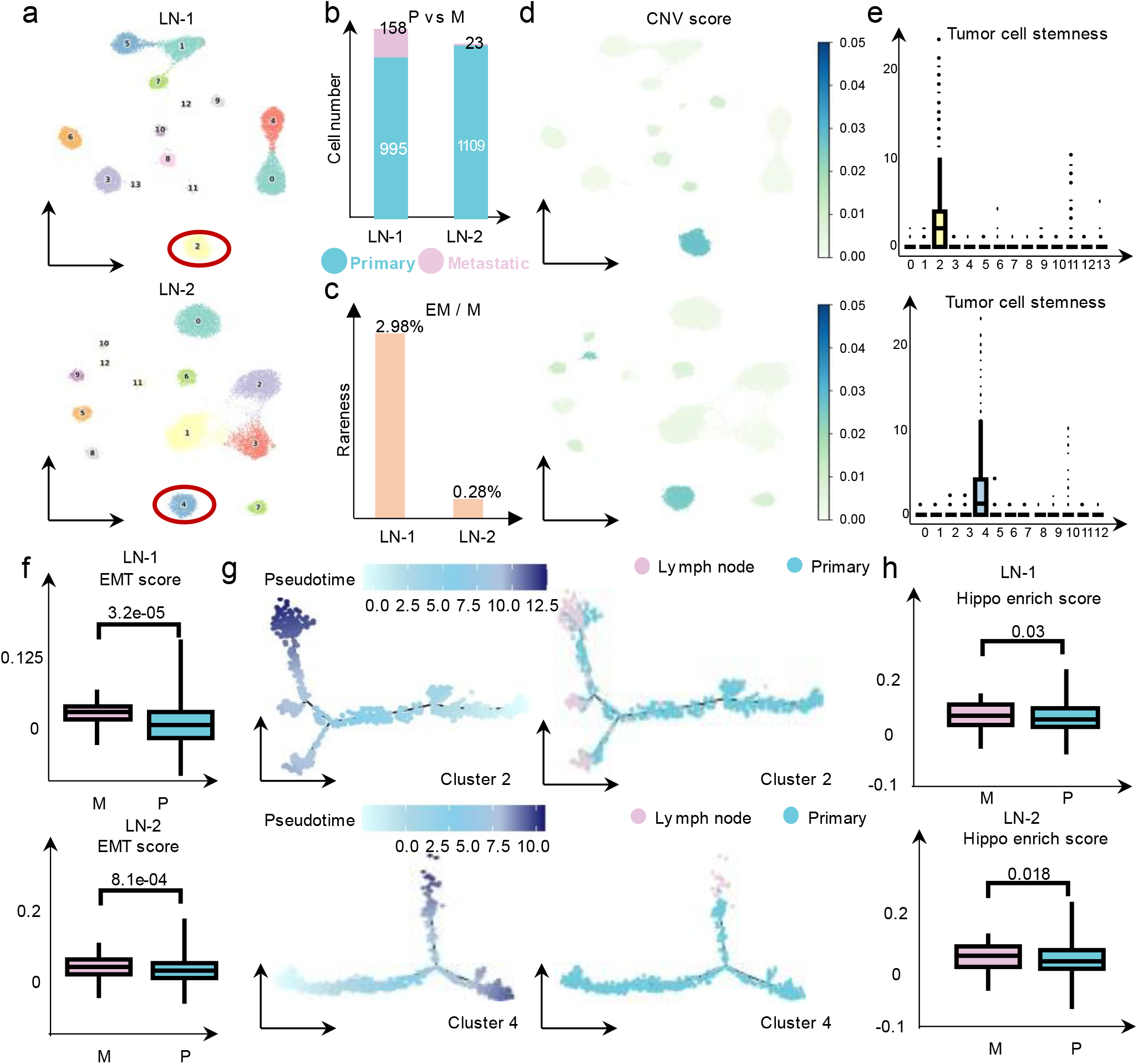
EmitGCL captures the occult metastatic cells in breast cancer metastasis patients with negative status lymph nodes. **a**, UMAP visualizes cell clusters predicted by EmitGCL in breast cancer with two negative lymph nodes (LN-1 and LN-2). The cluster with the red circle represents the metastatic precursor cell in primary tumor and the occult metastatic cells in the lymph node identified by EmitGCL. **b**, Bar plots show the cell number of the identified precursor (Blue) and occult metastatic cells (Pink) in LN-1 and LN-2. **c**, Bar plots show the proportion of occult metastatic cells with all cells in the lymph node. **d**, UMAP showcases the CNV score across all cells in LN-1 and LN-2. **e**, Box plots showcase the tumor cell stemness score across all cell clusters identified by EmitGCL in LN-1 and LN-2. **f**, Box plot showcases the EMT enrichment score in primary (P) and metastatic sites (M) across all cell clusters identified by EmitGCL in LN-1 and LN-2. Each box showcases the minimum, first quartile, median, third quartile, and maximum EMT enrichment scores on different sites (LN-1: P: = 995, M: = 158, LN-2: P: = 1109, M: = 23). The p-value is calculated by the Mann-Whitney U test with one-sided. **g**, The pseudotime of predicted precursor in primary tumor and associated metastatic cells in LN-1 and LN-2. Blue means the cell located in the primary, and pink means the cell located in the metastatic sites. **h**, Box plot showcases the metastasis-related pathway enrichment score in primary (P) and metastatic sites (M) across all cell clusters identified by EmitGCL in LN-1 and LN-2. Each box showcases the minimum, first quartile, median, third quartile, and maximum metastasis-related pathway enrichment scores on different sites (LN-1: P: N= 995, M: N= 158; LN-2: P: N= 1109, M: N= 23) N is cell number. The p-value is calculated by the Mann-Whitney U test with one-sided.

Given that metastatic cells could exhibit characteristics of later stages in cancer cell evolutionary progression^28^, we visualized the cellular trajectories and corresponding pseudotime for cluster 2 in LN-1 and cluster 4 in LN-2, respectively **(Fig. 3g and Supplementary Fig. 5a)**. This visualization further supports the identification of these clusters as true positives. Moreover, metastasis-related pathway enrichment scores were significantly higher in metastatic sites than in the primary site, consistent with the established knowledge that metastatic potential increases in cancer cells at metastatic sites. This may be driven in part by dysregulation of the Hippo signaling pathway, where increased YAP/TAZ activity promotes cancer cell survival, proliferation, and enhanced metastatic capacity^35-37^ **(Fig. 3h, Supplementary Fig. 5b and Supplementary Data 12)**. These findings provide strong evidence that EmitGCL can accurately identify occult metastatic cells that may be missed by conventional clinical methods, as well as metastatic precursor cells in primary tumors.

### Validation of biomarkers in precursor cells based on clinical cohort data

Previous validation demonstrated the superior performance of EmitGCL in identifying occult metastatic cells, providing a reliable foundation for metastatic precursor cell prediction. To identify biomarkers in metastatic precursor cells, we trained EmitGCL on data from 18 breast cancer patients. During model training, attention values were analyzed to identify biomarker genes specific to metastatic precursor and occult metastatic cells **(Methods) (Fig. 4a and Supplementary Data 13)**. Many of the genes identified in both metastatic precursor and occult metastatic cells, such as *ACTB*^*38*^, have already been validated as highly associated with breast cancer metastasis. The unique biomarkers in metastatic precursor cells identified from primary tumors were then used to build a regression model aimed at predicting future metastasis **(Methods)**.

**Fig. 4.**
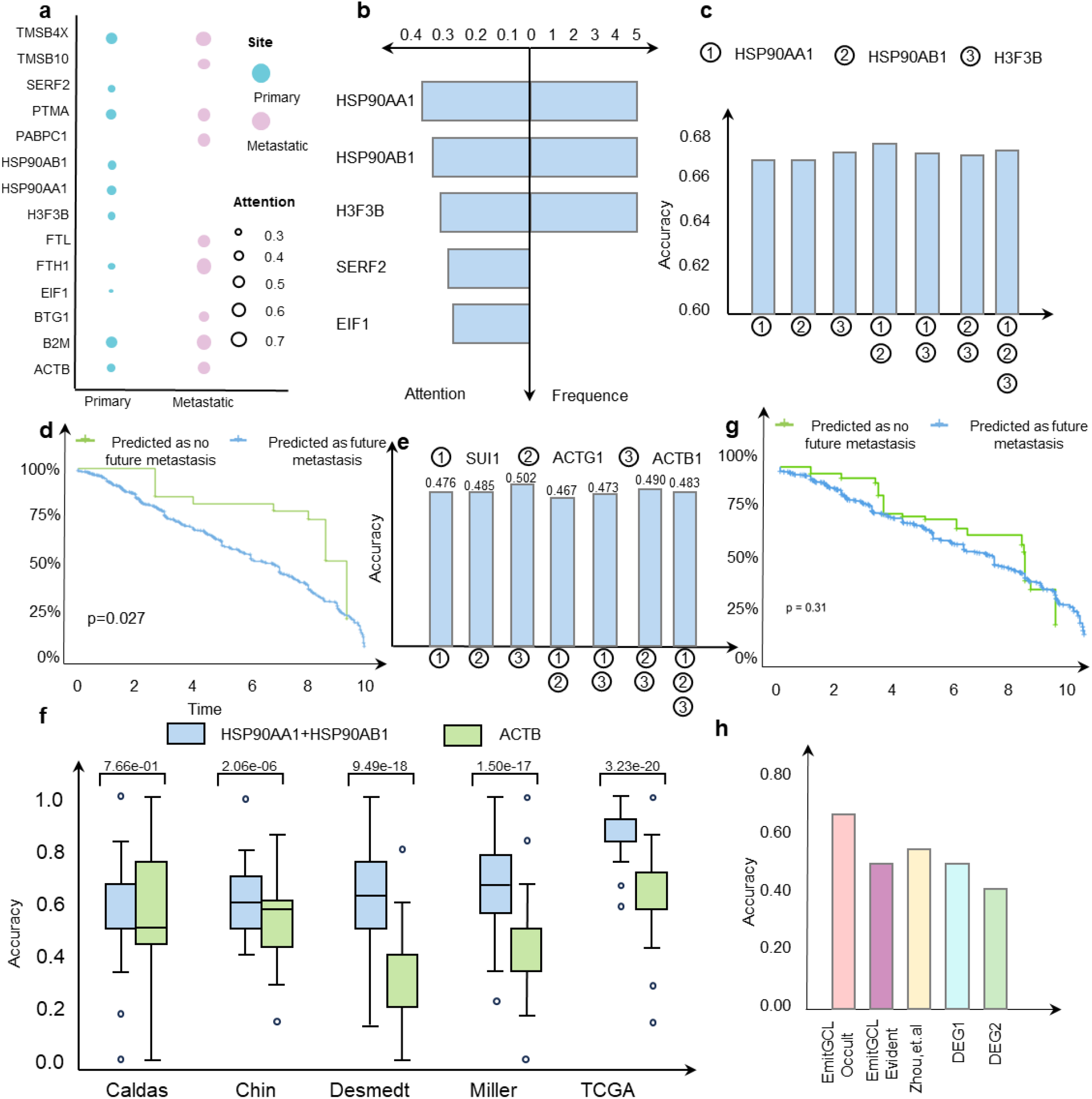
EmitGCL efficiently identifies breast cancer metastatic precursor gene biomarkers, demonstrating the potential for predicting the timing of occult metastasis in clinical settings. **a**, Dot plot showing the attention scores of gene biomarkers in the metastatic precursor (blue) cells in primary tumor and occult metastatic cells in lymph node. Colors represent different sites, and dot size indicates attention score. **b**, Bar plots showing the attention scores of gene biomarkers identified exclusively in metastatic precursor cells (left) and the frequency of these genes across five metastatic breast cancer cohorts (right). **c**, Bar plot showing the accuracy of predicting to develop metastasis and those without future metastasis using train/test dataset splits (9:1), based on biomarkers identified by tracing occult metastatic cells. **d**, Survival curve stratified by predicted future metastasis results using attention core of HSP90AA1 and HSP90AB1. **e**, Bar plot showing the accuracy of predicting to develop metastasis and those without future metastasis using a train/test dataset split (9:1), based on biomarkers identified by tracing clinically evident metastatic genes. **f**, Survival curve stratified by predicted future metastasis results using attention core of ACTB. **g**, Box plot showing the accuracy of predicting high versus low metastatic potential using a leave-one-dataset-out approach. Each box showcases the minimum, first quartile, median, third quartile, and maximum accuracy (Caldas: N=52, Chin: N=95; Desmedt: N=75; Miller: N=83, TCGA: N=115) N is the sample number. **h**, Bar plot comparing the performance for predicting biomarkers identified by, Z ‘ method, a cell clustering method based on DEGs (DEG1), and a cell classification method based on DEGs (DEG2).

Among the identified biomarkers, *HSP90AA1, HSP90AB1*, and *H3F3B* were consistently observed across five independent cohorts and exhibited high attention scores (**Fig. 4b)**. Notably, these biomarkers had contrasting fold-change values, rendering them undetectable through traditional differential expression analysis **(Supplementary Fig. 6a)**. To assess the accuracy and robustness of these biomarkers, we applied different train/test dataset splits and analyzed combinations of the three biomarkers. EmitGCL showed high accuracy across all splits, particularly in the 9:1 configuration, indicating robust performance. The combination of *HSP90AA1* and *HSP90AB1* delivered the best results and was selected for further analysis **(Fig. 4c, Supplementary Fig. 6b and Supplementary Data 14)**. A survival analysis based on future metastasis predictions using *HSP90AA1* and *HSP90AB1* revealed that patients who predicted future metastasis had significantly shorter overall survival compared to those without future metastasis **(Fig. 4d and Supplementary Data 15)**.

We hypothesize that metastatic precursor cells are better traced through occult metastatic cells rather than clinically evident metastatic cells. To validate this, we identified genes in metastatic precursor cells by tracing them through clinically evident metastatic cells. **(Supplementary Fig. 6c)**. The single gene *ACTB* yielded the best results and was selected for further analysis and comparison **(Fig. 4e)**. The validation using a leave-one-dataset-out approach across five cohorts also confirmed the consistent predictive accuracy of EmitGCL when trained on occult and clinically evident metastatic biomarkers (**Fig. 4e, f and Supplementary Data 16)**. Simultaneously, survival analysis using *ACTB* showed no significant difference in overall survival between patients predicted to develop metastasis and those without future metastasis **(Fig. 4g and Supplementary Data 17)**, confirming that EmitGCL based on clinically evident metastatic cells cannot reliably predict biomarkers of future metastasis.

To further validate the advantage of our method over the widely used DEG-based approaches, we examined DEGs between short-term and long-term metastases in each sample and found that no common DEGs were present across the five cohorts, and the large number of DEGs identified were unsuitable for clinical application due to their lack of specificity **(Supplementary Fig. 6d)**.

To provide an overall performance evaluation, we compared EmitGCL to other methods relying on DEGs. EmitGCL outperformed approaches based on clinically evident metastatic cells, as well as Zhoús TCGA based methods^6^, clustering-based (DEG1), and classification-based (DEG2) methods in predicting future metastasis **(Fig. 4h)**. In conclusion, this case study highlights the effectiveness of EmitGCL in identifying biomarkers of metastatic precursor cells and predicting future metastasis. This approach ultimately informs treatment strategies and improves patient prognosis.

### Validation of therapeutic target for preventing metastasis in-silico, experimentally, and clinically

Based on the genes identified in **(Fig. 4a)**, the gene regulatory networks in both sites were inferred using the ChEA3 database^39^. The importance of TFs was ranked across multiple metrics, and the high-ranked TFs will undergo simulation and experimental validation **(Supplementary Fig. 7a)**. The resulting regulatory networks in primary and metastatic sites are displayed in **Fig. 5a**. We evaluated all regulators and regulons based on TF frequency, TF expression, and regulon attention scores, identifying 12 regulons with high scores across all indices **(Fig. 5b and Supplementary Data 18)**. Among them, JUN^40^ and FOS^41^ had the highest scores and have been validated as highly related to metastasis in previous studies. To further validate TFs that have not yet been confirmed in existing publications, we selected YY1, which demonstrated high TF closeness within the metastatic regulatory networks and is also a component of the polycomb repressive complex (PRC). PRC functions as an enhancer, promoting oncogenic transcriptional programs^42^. Another PRC component, EZH2, has been shown to drive breast cancer metastasis β -FAK activation^43^. However, the role of YY1 in this process has not been previously investigated.

We initially conducted *in-silico* perturbation using CellOracle^24^ for each breast cancer patient. The cell clusters identified by EmitGCL for each patient were visualized using UMAP, with cluster annotations determined based on the marker genes corresponding to each cell type **(Supplementary Fig. 7b and Supplementary Fig. 8a)**. Based on the identified cell clusters, epithelial and cancer cells were extracted as inputs for simulated knockout validation using CellOracle. The results revealed that cells with high metastatic potential tended to transition toward normal epithelial cells, supporting the hypothesis that YY1 serves critical TF in developing metastasis **(Fig. 5c and Supplementary Fig. 8b)**.

**Fig. 5.**
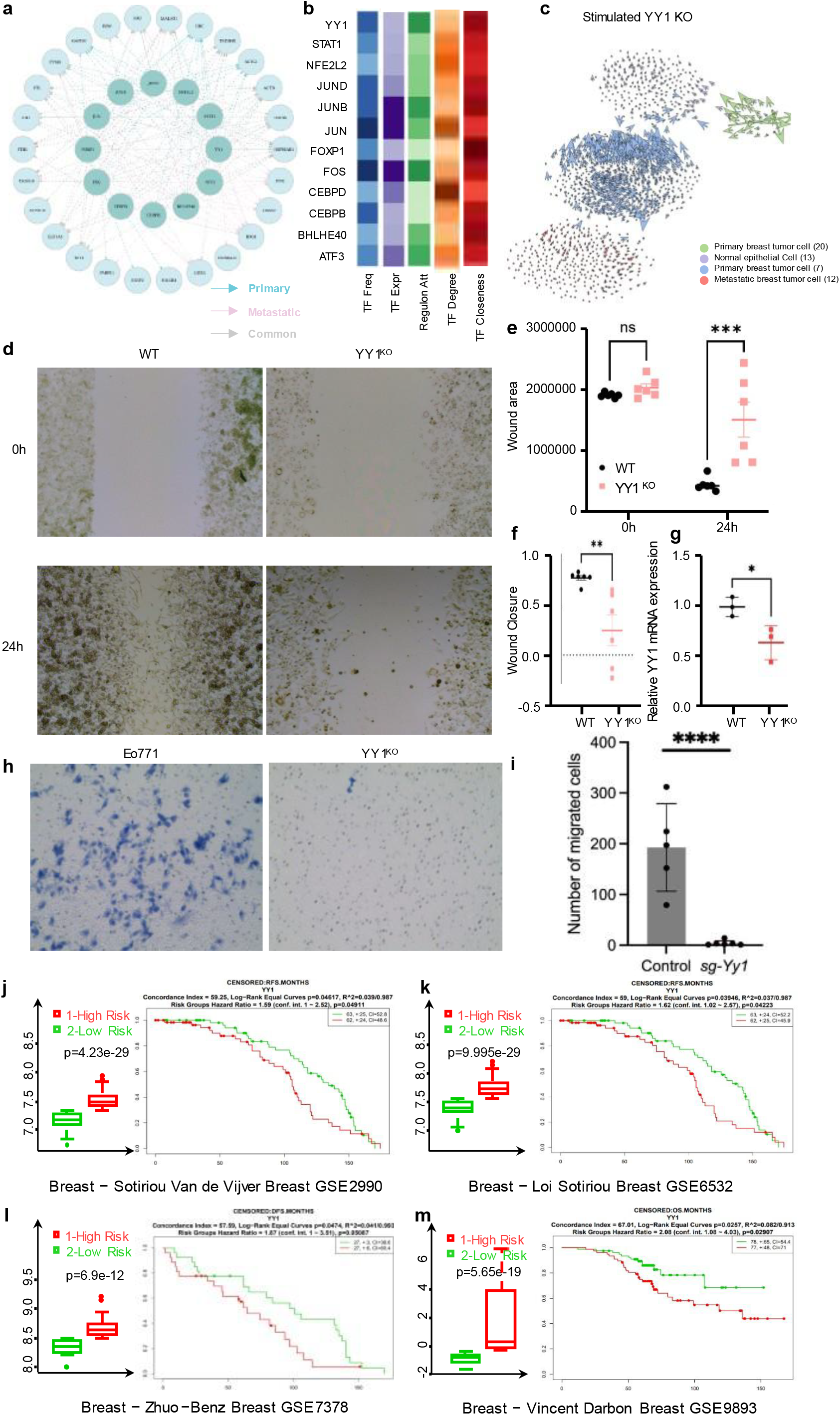
EmitGCL experimentally and clinically validates critical TFs for inducing breast cancer metastasis. **a**, TF regulatory networks of precursor and occult metastatic cells across all breast cancer patients. Blue indicates regulatory relationships in precursor cells, pink in occult metastatic cells, and gray for common regulatory relationships. **b**, Heatmap TF frequency, TF expression, regulon activity score, degree and closeness centrality scores for the regulatory network shown in panel b. **c**, Observed and extrapolated future states (arrows) following YY1 knockout in four epithelial subpopulations. Colors represent different cell clusters. **d**, The wound healing assay with YY1^KO^ cells at 24 hours compared to the control group. **e**, The wound healing areas in YY1^KO^ cell and control group (**p<0.01, ***p<0.001). **f**, The migration capacity of YY1^KO^ cell and control group. **g**, The total RNA was isolated from Control and Yy ^KO^ group, and relative YY_1_ mRNA expression was measured by qPCR. Data means t SD (n=3). p < 0.05, indicating significant reduction in YY1 expression in YY1^KO^ compared to Control. **h**, Representative images of the transwell migration assay for parenteral EO771 cells and it with CRISPR-mediated knockout of YY1. Migrated cells were stained with 4% Trypan Blue. **i**, Quantification of Invasive Cells by light microscopic evaluation. Each group had six independent replicates. The number of invading cells was compared between parenteral EO771 cells and CRISPR-mediated knockout of YY1 (****p<0.0001). j, The box plots represent the stratification of patients based on the median expression level of YY1 in GSE2990 cohort. The Kaplan-Meier curves depict relapse-free survival outcomes, stratified by high and low YY1. **k**, The box plots represent the stratification of patients based on the median expression level of YY1 in GSE6532 cohort. The Kaplan curves depict relapse-free survival outcomes, stratified by high and low YY1. **l**, The box plots represent the stratification of patients based on the median expression level of YY1 in GSE6532 cohort. The Kaplan curves depict relapse-free survival outcomes, stratified by high and low YY1. **m**, The box plots represent the stratification of patients based on the median expression level of YY1 in GSE6532 cohort. The Kaplan curves depict overall survival outcomes, stratified by high and low YY1.

To further validate the role of key TFs in the occult metastasis process, we first used CRISPR-mediated knockout of FOXP1 and JUND in EO771 cells and demonstrated a significant deficiency in cell migratory capacity by transwell cell migration assay (**Supplementary Fig. 9c and Supplementary Data 19)**. This finding aligns with previous studies indicating the pivotal role of these TFs in cancer metastasis. Subsequently, we focused on the newly identified TF, YY1, and observed that YY1^KO^ cells exhibit reduced wound closure at 24 hours compared to the control group **(Fig. 5d and Supplementary Fig. 10 and Supplementary Data 19)**. Quantification of wound healing areas and wound closure speed revealed a significant reduction in YY1^KO^ cells compared to control groups (**p<0.01, ***p<0.001) **(Fig. 5e and 5f)**. YY1 mRNA expression measured by qPCR indicated a significant reduction in YY1 expression in YY1^KO^ compared to Control (*p<0.05) **(Fig. 5g)**. To more accurately investigate the ability of tumor cells to invade across the basement membrane, rather than merely assessing lateral migratory or proliferative capacity as in wound healing assays. We performed a transwell cell migration assay and revealed that YY1^KO^ cells had a significant reduction in cell migratory capacity compared to the control group (****p<0.0001) **(Fig. 5h and 5i)**.

To further validate the role of YY1 in the survival outcomes of breast cancer patients, four independent cohort studies with bulk RNA sequencing data were analyzed. Patients in each cohort were stratified into high and low YY1 expression groups based on the median expression level. Relapse-free survival was assessed in cohorts 1 to 3, while overall survival was evaluated in cohort 4, using Kaplan-Meier curves for all analyses. Survival differences were compared using the log-rank test. In cohorts 1 to 3, patients with YY1 expression levels higher than the median exhibited significantly shorter relapse-free survival, with hazard ratios of 1.59 [95% confidence interval (CI), 1.00–2.52] (p=0.049), 1.62 [95% CI, 1.02–2.57] (p=0.042), and 1.87 [95% CI, 1.00– 3.51] (p=0.050) **(Fig. 5j-l)**, respectively. Similarly, patients with higher YY1 expression in cohort 4 demonstrated significantly shorter overall survival, with a hazard ratio of 2.08 [95% CI, 1.08–4.03] (p=0.029) **(Fig. 5m)**.

### EmitGCL reveals a more immunosuppressive environment in lymph node sites compared to primary sites in breast cancer

To understand the differences between primary and metastatic sites during the metastatic process, we performed pathway enrichment analysis on gene signatures from both locations **(Fig. 6a)**. The results revealed that immune-related pathways, such as antigen processing and presentation, were more significantly enriched in the primary site, suggesting a more active immune environment compared to the metastatic site. To explore the underlying reasons for this observation, we used CellChat to analyze cell-cell communication between early metastatic cell groups and cytotoxic T cells, as well as between early metastatic cell groups and antigen-presenting cells. Our analysis showed that the MIF signaling pathway, known for its immunosuppressive effects, had a higher communication probability between early metastatic cell groups and cytotoxic T cells compared to precursor cell groups and cytotoxic T cells **(Fig. 6b)**. Further, we calculated the T cell exhaustion and effective activity scores at both primary and lymph node sites. The results revealed that T cells exhibited higher exhaustion scores in lymph nodes, indicating a more immunosuppressive environment, while showing higher effective activity in primary sites **(Fig. 6c)**. This was corroborated by the bar plot of T cell clonotypes, where lymph nodes demonstrated more clonotype expansion than primary sites, highlighting the progression of immune suppression in metastatic environments **(Fig. 6d-f)**. Additionally, pathway enrichment analysis based on differentially expressed genes (DEGs) between precursor and metastatic cell groups further supported the presence of enhanced immunosuppressive mechanisms in the lymph node sites **(Fig. 6g)**.

**Fig. 6.**
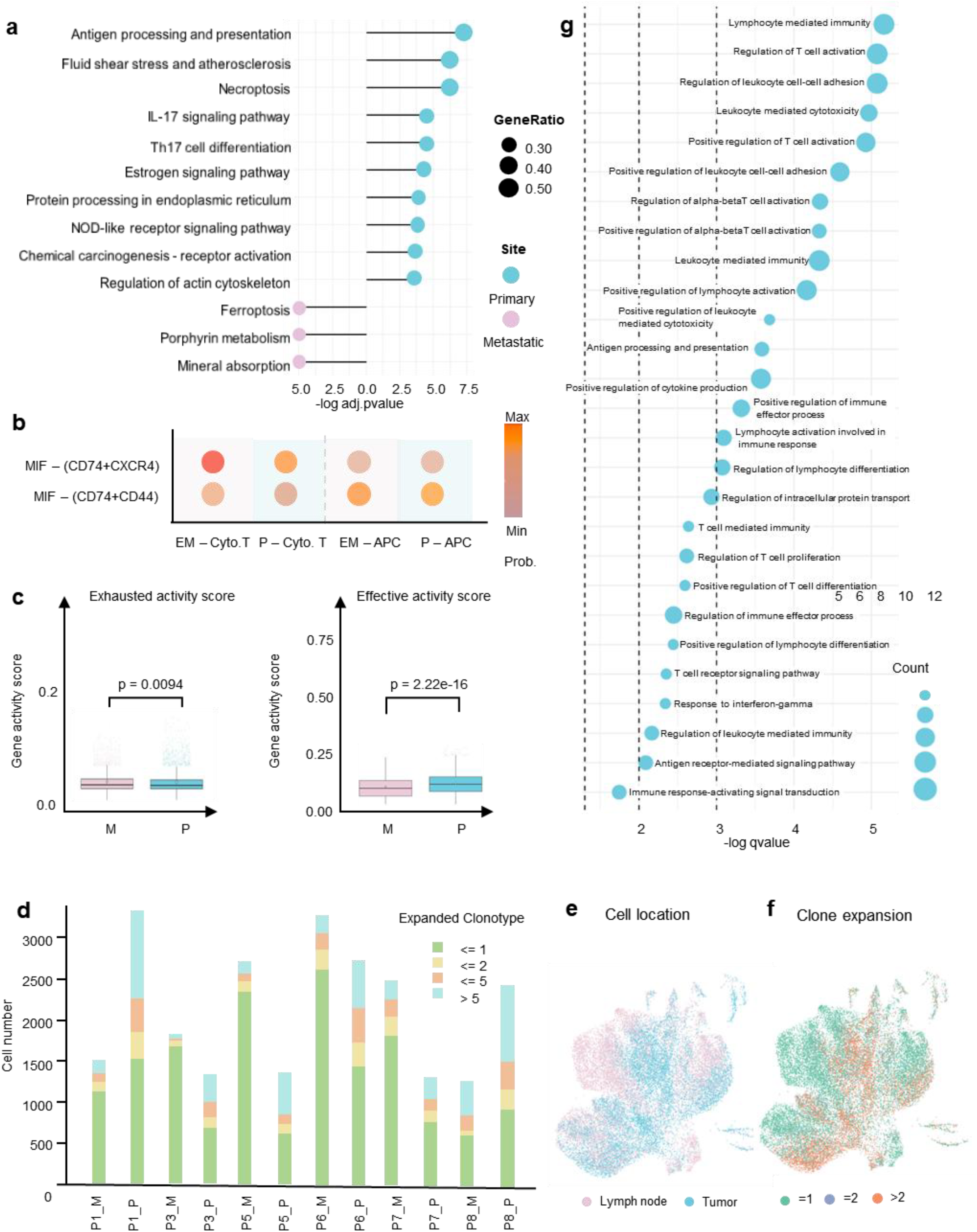
EmitGCL reveals a more immunosuppressive environment in lymph node sites compared to primary sites in breast cancer. **a**, Pathway enrichment analysis across precursor (blue) and early metastatic cell groups (pink). Colors represent different sites, and dot size indicates the ratio of enriched genes. p-values were calculated using a one-sided hypergeometric test, with FDR-adjusted p-values for multiple testing correction. **b**, MIF signaling pathway analysis of cell-cell communication between precursor and early metastatic cell groups and cytotoxic/antigen-presenting cells. Colors represent communication probability. **c**, Enrichment scores of effective and exhausted gene signatures across different sites in cytotoxic T cells (Primary: ***n*** = 176, Metastasis: ***n*** = 895). Box plots display the range of gene signature enrichment scores. p-values were calculated using the two-sided Mann-Whitney U test. **d**, Bar plot showing the number of T clonotypes in primary and metastatic sites. e, Cell location across all samples based on scTCR-seq data. UMAP plot showing the distribution of cells between primary tumor (green) and lymph node metastasis sites (blue). **f**, Clone expansion across primary and lymph node sites. UMAP plot highlighting T cell clonal expansion, with color intensity (red) indicating higher clonal expansion, showing greater expansion in lymph node sites compared to primary sites. **g**, Immune-related pathway enrichment based on DEGs between precursor and metastatic cell groups. Dot size indicates the count of enriched genes. q-values were calculated using a one-sided hypergeometric test with multiple testing correction.

## Discussion

EmitGCL represents a significant advancement in predicting future metastasis based on scRNA-seq, particularly in complex biological settings where traditional computational methods and clinical diagnostic technologies often fall short. By integrating prior knowledge of metastasis, EmitGCL achieves high TPR while maintaining a low FPR in occult metastatic cell identification. This accuracy provides the necessary foundation for reliable metastatic precursor cell inference. Leveraging graph contrastive learning, EmitGCL enables the tracing of dynamic metastatic transitions between metastatic precursor cells and occult metastatic cells.

In this study, which involved 28 patients and 516,093 cells across six cancer types, we demonstrated the exceptional performance of EmitGCL in identifying occult metastatic cells through computational validation. While direct targeted attempts for the HSP90 complex have not been successful clinically, the HSP90 complex is highly critical to the integrity and function of many oncogenes by maintaining their protein folding and stability.These genes being primary markers for metastatic precursor cells aligns with the knowledge that the process of metastasis is highly stressful to cancer cells and the cellular response to this stress and maintenance of protein integrity are likely key to successful metastasis. However, these genes could prove to be good biomarkers for the likelihood of metastasis and delineate whether primary tumors need additional treatment to prevent future metastasis from occurring. Additionally, EmitGCL uncovered the key transcription factor YY1, which plays a role in driving breast cancer metastasis. YY1 has been shown to play various roles in tumors, including regulating expression of oncogenes and tumor suppressor genes, metabolic reprogramming, and promoting the cancer stem cell phenotype. YY1 has also been shown to regulate the tumor microenvironment by promoting angiogenesis and suppressing immune-mediated killing, among others. These functions accomplished by YY1 would be highly beneficial and likely necessary for metastasis to be successful. YY1 was likely detected by EmitGCL for these reasons and will provide promise for enhancing therapeutic strategies and providing insights for future clinical trial design.

Despite these promising results, some limitations of EmitGCL remain. First, a major limitation is the lack of perspective clinical trial validation for the biomarkers identified by EmitGCL. Without rigorous clinical testing, the translational potential of these findings remains speculative. Future studies will focus on validating these biomarkers in clinical trials to ensure their robustness ‘ into actionable strategies in precision medicine, ultimately improving outcomes for patients at high risk of metastasis. Second, the current predictive model was developed based on data from seven cohorts comprising 28 patients, which exhibited considerable heterogeneity in clinical behavior. Future large-scale studies with a greater number of patients are needed to validate these findings and improve model generalizability.

In conclusion, EmitGCL offers a prior knowledge-aware graph contrastive learning model capable of predicting future metastasis and their biomarkers. It also uncovers key TFs that can be regarded as therapeutic targets in clinical applications. EmitGCL provides valuable insights that could inform potential therapeutic strategies and clinical trial designs, ultimately holding promise for improving patient outcomes.

## Methods

### Cell culture

EO771 cells were maintained in Dulbeccós 2 mM glutamine, 100 U ml^−1^ penicillin, 100 μg ml^−1^ streptomycin and 10% fetal calf serum (FCS).

### CRISPR knockout

CRISPR/Cas9 system was used to knock out the gene encoding transcription factor *Yy1, Foxp1*, and *Jund* in EO771 cell line via electroporation. Two predesigned sgRNAs are selected to target each gene: *Yy1* ′-GCCCACCACCGTGGTCTCGA- ‘ ′-ACCCTCTACATCGCCACGGA- ‘, *Foxp1* ‘-CTTCGTGACACTCGGTCCAA- ‘, ‘-TAGTAAGTGGTTGCCACCGC- ‘,, *Jund* ‘- TACGCAGTTCCTCTACCCGA- ‘, ‘-GATCATCCAGTCCAACGGGC- ‘, Cas9 Nuclease and electroporation enhancer were acquired from IDT. Electroporation was performed following the protocol provided by IDT CRISPR genome editing (www.idtdna.com) using the Lonza 4D Nucleofector System.

### Transwell migration assay

EO771 cells were subjected to CRISPR-based gene editing to generate knockout groups. Following transfection, cells were cultured for 48 hours. Each group was then diluted to a concentration of 20,000 cells in 200ul 10% fetal bovine serum (FBS)-containing medium and seeded into the upper chamber of transwell inserts. The lower chambers were filled with 600 μL of 10% FBS-containing medium. After incubation for approximately 36 hours, inserts were collected, and non-invading cells on the upper surface of the membrane were gently removed using a cotton swab. The invaded cells were fixed in 70% ethanol for 1 minute, followed by staining with 4% Trypan Blue for 10 minutes. The inserts were then washed twice with PBS. Images of invaded cells were captured using a light microscope for further analysis.

### Data description

We included 33 publicly available matched scRNA-seq datasets from primary and metastatic sites across 28 patients with six cancer types in the paper. All cells were used for model training. The paired lung and two lymph node scRNA-seq datasets from a lung cancer patient without metastasis were used for false positive evaluation in the benchmark section. The paired pancreatic and liver scRNA-seq data from pancreatic cancer patients with liver metastasis, along with data from four other cancer types (breast, nasopharyngeal, papillary thyroid, and head and neck), which exhibited lymph node metastasis, were used for sensitivity testing in the benchmark section. The paired breast and lymph node scRNA-seq datasets from 13 breast cancer patients with metastasis were used for case analysis. Additionally, 420 bulk breast cancer RNA-seq datasets from UCSC were used to validate the biomarkers identified by EmitGCL. All data are publicly available (**Supplementary Data 1**).

### Model’s inputs

EmitGCL initiates by inputting two count matrices 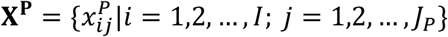 and 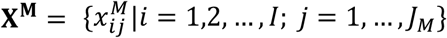 derived from matched scRNA-seq data from primary and metastatic sites. We organize the data such that rows represent genes, whereas cells constitute the columns. Any row or column in each count matrix that contains only all zero values will be excluded from further analysis.

### Model’s inputs

After the model training, we can obtain results at two levels. At the cell level, we identified: (*i*) occult metastatic cells in metastatic sites, (*ii*) metastatic precursor cells which are defined as the cells that are predicted in the same cluster as the occult metastatic cell we identified in (*i*). At gene level, we identified biomarkers of metastatic precursor cells for future metastasis prediction.

### Heterogeneous bipartite graph construction

To formulate the cross-modal relationships between primary and metastatic site, the matched scRNA-seq data is represented by one heterogeneous bipartite graph, comprising nodes of cells, genes, which captures these interactions and dependencies into a unified framework.

In our case, we integrate matrices **X**^**P**^ and **X**^**M**^ into a combined matrix **X** = [**X**^*P*^, **X**^*M*^] by constructing a gene-cell heterogeneous bipartite graph **G**, consisting of two node types and one edge type to ensure each type of element (nodes and edges) maintains a unique distribution and furnishes a natural representation framework. We define the heterogeneous graph as **G** = (**V, S, E, F**) with node set **V** = **V**^**C**^ ∪ **V**^**G**^, where 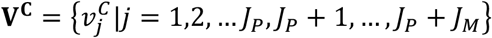 denotes all cells, 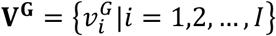 denotes all genes. The sign set is a binary vector **S** = {*s*_*j*_|*s*_*j*_ ∈ {0,1}^*J*^}. *s*^*j*^ = 0 means the cell *s*^*j*^ is in primary sites, and *s*^*j*^ = 1 means the cell is in metastatic sites. The edge set **E** is constituted as 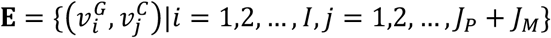, with edge weight *w* defined as follows. To eliminate information redundancy between node initial embeddings and the edge weights, we utilize unweighted edges when constructing the heterogeneous bipartite graph. For 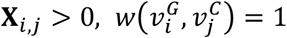, otherwise, 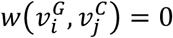. Lastly, we establish the initial feature vectors **F** for nodes in *G* as follows:

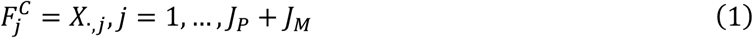

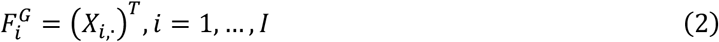

where *X*_*i*,._ and *X*_.,*j*_ represent the *i*_*th*_ row vector and the *j*_*th*_ column vector of **X**, respectively.

### Sub-sampling of a heterogeneous bipartite graph

To enhance the efficiency of EmitGCL when dealing with a large heterogeneous graph, it is necessary to select subgraphs before model training. Considering the rarity of occult metastatic cells in metastatic sites, no such requirement exists in the primary site. We select the subgraph differently for each site. In the primary site, for a targeted cell 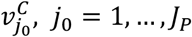, the genes 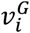 linked to the targeted cell 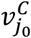 are randomly selected. We consider that a gene ubiquitously expressed is unlikely to hold as much significance for rare cell identification compared to a gene that is expressed only in a particular subpopulation. Therefore, we devised a probability-based sub-sampling method in metastatic sites to select the subgraphs. There are two steps for the probability-based sub-sampling method. The first step is to filter out low expressed genes, which should not be regarded as rare-related features. For a targeted cell 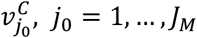, the genes 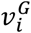, linked to the targeted cell 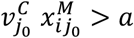 are reserved. *a* is a threshold set to the first quartile of the expression value of all genes in the given cell. The second step is to select genes 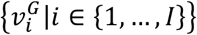 based on probabilities calculated by the following formula:

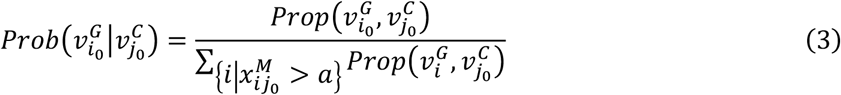

where

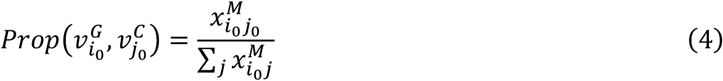

The higher the probability value of gene 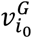 correspondence to cell 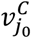, the more likely it is to be selected into the subgraph for cell 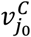. At this point, the gene has a higher probability of displaying the cell’s rare signal. The number of reserved genes is denoted as *N*_*g*_. Considering the computational expense of the deep learning model, we set the number of selected genes to min( *N*_*g*_, *nbatch*) by default. The *nbatch* is a hyperparameter which means batch number. In each site, the subgraph incorporates 30 randomly selected cells along with their chosen neighbor nodes. EmitGCL is trained using multiple mini batches, each represented by a subgraph.

### EmitGCL’s embedding updated by transformer

Considering the importance of both local and global information for embedding updates, we employ a graph transformer model to facilitate message passing between cells and genes. This model effectively mitigates the impact of missing values and dropout issues commonly encountered in single-cell data by leveraging the connectivity and relationships within the graph to enhance the robustness and accuracy of the embeddings.

Let **H**^***l***^ represent the embedding of the *l*^*th*^ layer (*l* =1,2, …, *L*). The updated embedding of 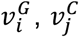, on the *l*^*th*^ layer is denoted as 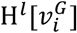 and 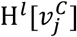, respectively. To align the features of different types of node type into the same dimension, we apply a linear projection function *W*^*G*^ on the gene 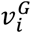 initial feature vectors ***F***^*G*^ to obtain the initial embeddings 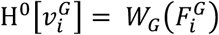. We also apply a linear projection function *W*^*C*^ on the cell 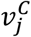 initial feature vectors ***F***^*C*^ to obtain the initial embeddings 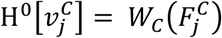. Both embedding dimensions are set to the same lower value *d*.

To enable the model, capture a broader range of dependencies and patterns by focusing on different parts of the input simultaneously, thereby improving representation power and mitigating information loss. We use a multi-head mechanism, dividing H^0^[𝓋] into **H** heads. For the *h*^*th*^ attention head in the *l*^*th*^ layer, the attention value between node 𝓋 and the one-hop neighbor 𝓋^*ne*^ is calculated as follows:

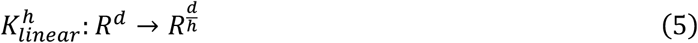

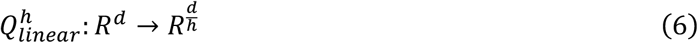

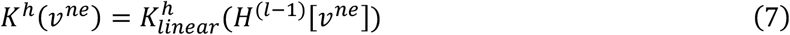

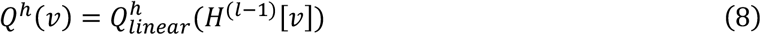

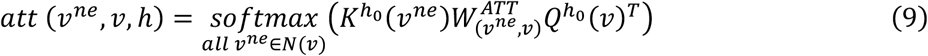

where *K*^*h*^ and *Q*^*h*^ are two different linear projection functions, 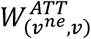 is a transformation matrix designed to capture edge features, (.)^*T*^ signifies the transposal function. *N*(𝓋) is the neighbor node set of 𝓋.

The concatenation of attention heads yields the attention coefficients, represented as follows:

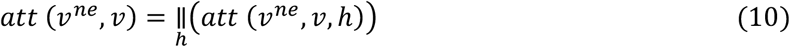

The message from 𝓋^*ne*^ that can be relayed to 𝓋 within head *h* is given by 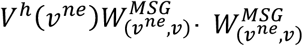 is also a transformation matrix and *V*^*h*^ is also a linear projection function. Then, the results from different message heads should subsequently be concatenated:

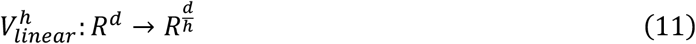

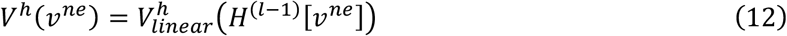

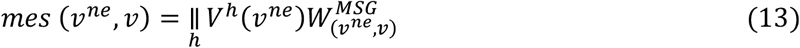

To update the embedding of node 𝓋, the final step within the *l*^*th*^ layer will sum *H*^*l*−1^[𝓋] and *H*^*l*′^[𝓋] with trainable weights into the node’s new embedding.

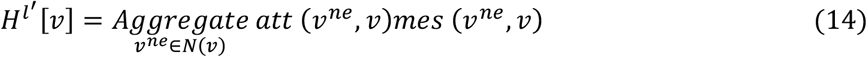

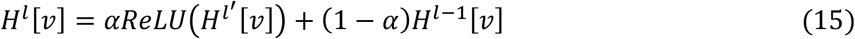

where α represents a trainable parameter, while ReLU functions as the activation function. The final embedding of 𝓋 is obtained by layer-wise stacking of information.

### EmitGCL subgraph training

In the context of handling large and complex graphs, memory usage and computational efficiency are significant concerns. EmitGCL addresses these issues by implementing subgraph training, which involves focusing on smaller, more manageable portions of the graph. This approach significantly reduces memory requirements and computational costs, making it possible to process and analyze large-scale graphs more effectively. By concentrating resources on these subgraphs, EmitGCL can operate more efficiently, allowing for faster training times and lower memory consumption, while still capturing the essential information needed for accurate modeling.

### EmitGCL loss design

In our model training, we devise a multi-task loss function, which consists of four critical components. Loss component (1) is designed to obtain high-quality node embeddings. To reduce the false positive in the occult metastatic cells we identified, we design loss component (2) to involve prior biological information, the metastasis-related pathways, to penalize the rare cell we identified as occult metastatic cells but with low metastasis-related active score. Then, to improve the sensitivity of the occult metastatic cell identification, we design the loss component (3), a contrastive learning loss to enhance the difference between the primary site and metastatic sites, which helps us to capture slightly changes and the metastatic trail. The loss component (4) is the key to keeping the balance of the rare and major cell group signal. The formulation of the multi-task loss function is defined as follows:

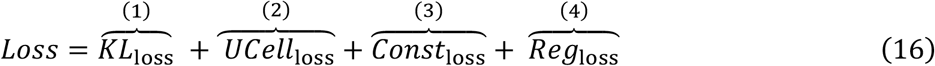

Where 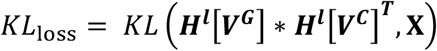. We consider occult metastatic cells as rare cell groups in the metastatic sites. Due to this property, false positives may occur. To reduce false positives, we incorporate prior knowledge of metastatic-related pathways. We then calculate the activation score of these pathways in the rare cell groups identified in the metastatic sites. The *UCell*^loss^ is defined as:

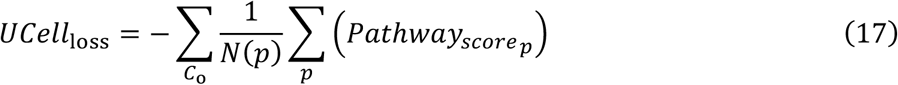

Where *C*^o^ is a cell set which is identified as the candidate occult metastatic cells. *N*(*C*^o^) is the cell number of *C*^o^, *p* is the metastasis-related pathway we used as the prior biological information to reduce false positive of prediction. *N*(*p*) is the pathway number we used. The 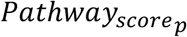 is the pathway *p* active score, which is defined as:

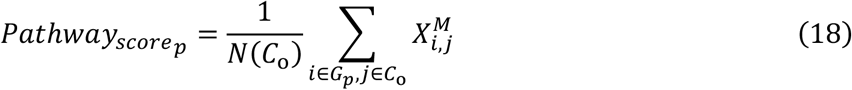

Where *G* ^*p*^ is a gene set in pathway *p*.

To depict the metastatic trail from the primary site to metastatic sites and enhance the model’ sensitivity, we designed the contrastive loss *Const*^loss^ with three distance comparisons:

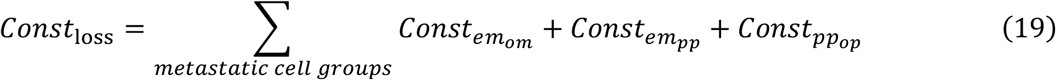

First, consider that the distance between cells in the same occult metastatic cells should be shorter than the distance between occult metastatic cells and other cells in the metastatic sites. We set the distance between cells in the same occult metastatic cells as the positive sample pairs and the distance between occult metastatic cells and other cells in the metastatic sites as the negative sample pairs. This involves the contrastive loss term 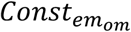 which is defined as follows:

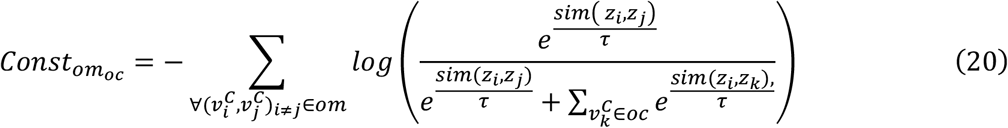

Where *om* is the identified occult metastatic cell set, *oc* is set of other cells in metastatic sites. *z*_*i*_, *z*_*j*_, *z*_*k*_ represents the embedding of cell 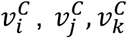, T concentration level of the distribution. Lower values make the model focus more on the most similar pairs, while higher values spread the focus over a larger set of pairs. *log* (*⋅*) is the nature logarithm function, which is applied to the probability to ensure numerical stability and to bring the loss value to a suitable range. *e*^*⋅*^ means the exponential function, which is applied to the similar scores divided by the temperature to convert them into a probability-like value. *sim*( *z*^*i*^, *z*^*j*^) means the similarity between cell 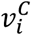 and cell 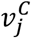 which defined as:

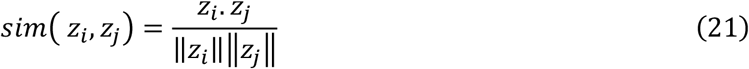

Where *⋅* denotes the dot product and || *⋅* || denotes the *L*_2_ norm.

To enhance the difference between the metastatic precursor cells and the occult metastatic cells, we designed the contrastive loss 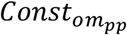 as follows:

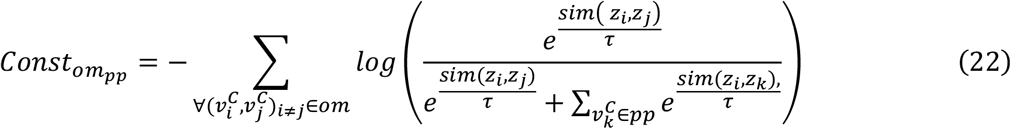

Where *om* is the predicted occult metastatic cell set, *pp* is the precursor cell set.

Furthermore, consider the difference between the precursor cells and other cells in primary site, we design the third contrastive loss:

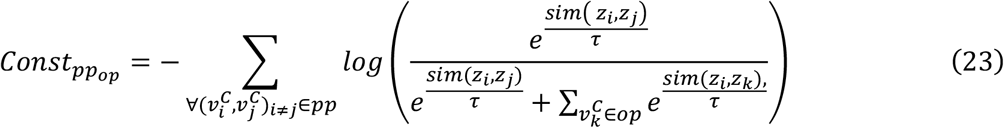

Where *pp* is the predicted occult metastatic cell set, *op* is set of other cells in primary site. Combining the three contrastive learning losses allows us to more sensitively identify occult metastatic cells and precursor cells, thereby depicting the tumor metastasis process.

Finally, to avoid losing all major cell information in metastatic sites, we designed a regularized term *Reg*^loss^ as follows:

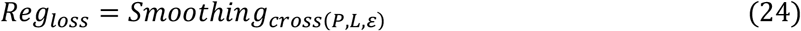

where *L* = {*l*_*j*_|*l*_*j*_ ∈ {1, … , *T*}, *j* = 1, … , *J*_*P*_ + *J*_*M*_} is the cell cluster results by Louvain with scRNA-seq from both primary and metastatic sites, *P* is the predicted cell cluster results of the model. *T* is the number of cell clusters by Louvain, ε is a smoothing factor.

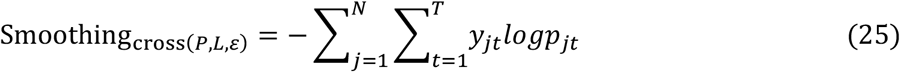

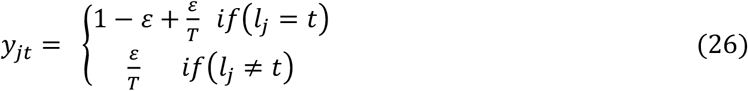

where *p*^*jt*^ represents the predicted probability of the given cell 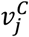 belonging to the class *t*.

### EmitGCL identified occult metastatic and precursor cells

After the training finished, we obtained the cell probability matrix *P*_*J*×*F*_ . The row represents cell, the column represents predicted cell groups. The value in *P*_*J*×*F*_ means the probability of 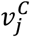 belongs to cell group *f*. The cells will be defined as occult metastatic cells if (*i*) in metastatic site, (*ii*) the cell number in the candidate cell group is less than 3%, (*iii*) the cell group with high tumor marker expression value, (*iv*) The cell number of metastatic site in candidate cell groups is lower than that in the primary site.

The cells will be defined as precursor cells if the cells are in the same cell group with identified occult metastatic cells.

### EmitGCL identified biomarkers

After the training, we obtained all cell groups of the primary and metastatic sites. For a target cell 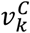 , we can obtain an attention matrix *A* = {*A*_*i*,*k*,*h*_|*i* = 1,2, … , *I, h* = 1, 2, … , *H*}, where the rows represent genes, and the columns represent head. The elements in the attention matrix indicate the contribution of each gene to the target cell in the cell clustering results. Considering that both negative and positive attention can contribute to distinguishing cell clusters, we incorporate the attention values across all heads in the target cell using an absolute value operation. If a gene has a high attention value across all heads, it will be identified as a gene signature of the target cell. For a targeted cell group *T* we identified, the contribution vector *C*_*T*_ with *I* length represents the contribution of all genes to the targeted cell group *T* which is defined as:

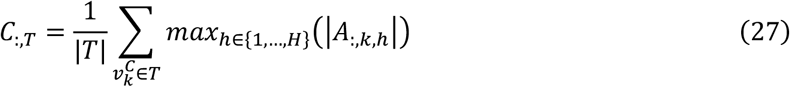

Then, we get a contribution matrix *C* of all genes to all cluster groups. The row of the matrix represents the gene, the column represents the cell group we identified in our model.

### Baseline tools parameter set

To evaluate the performance of EmitGCL relative to other tools for the FPR and sensitivity in occult metastatic cell identification, we conducted a comparative analysis between EmitGCL and other established methods.

1. **MarsGT**^**25**^ (Python package, v 0.2.1, https://github.com/OSU-BMBL/marsgt): Our in-house tool for rare cell identification, which demonstrated the best performance overall. Both the dimensionality reduction function “NodeDimensionReduction” and the prediction function “MarsGT_pred” were executed using default parameters.
2. **CellSIUS**^**27**^ (R package, v 1.0.0, https://github.com/Novartis/CellSIUS): Cells were filtered based on the total number of detected genes, total UMI counts, and the percentage of total UMI counts attributed to mitochondrial genes. Genes had to be present with at least 3 UMIs in at least one cell. After this initial QC, the remaining outlier cells were identified and removed using the “plotPCA” function from the scatter package (with detect_outliers set to TRUE). Data were normalized using the scran package, including a first clustering step as implemented in the “quickCluster” function.
3. **GiniClust**^**17**^ (Python package, v 3.0, https://github.com/rdong08/GiniClust3 : The “neighbors” parameter of the function “clusterGini” was set to 10 (the recommended value is from 5 to 15), while other parameters were kept at their default values.
4. **Seurat**^**26**^ (R package, v 4.3.0, https://satijalab.org/seurat : The “FindClusters” function was used for clustering with default parameters. Other functions, such as “NormalizeData”, “FindVariableFeatures”, “RunPCA”, and “RunUMAP”, were also executed using default settings.

### Rareness

Defined as the proportion of occult metastatic cells relative to the total number of cells in metastatic sites.

### Gene activity score

We used the AddModuleScore_Ucell function from the UCell package^44^ (R package, v 2.3.1) to calculate gene activity scores for different reference gene sets. This function applies the Wilcoxon rank-sum test (U statistic) to assess the activity of gene sets by comparing the expression levels of each gene within the set to all other genes. The output is a relative activity score computed for each cell.

### EMT activity score

To calculate the EMT activity score, we retrieved the HALLMARK_EPITHELIAL_MESENCHYMAL_TRANSITION gene set from the MSigDB database using the msigdbr package (R package, v 7.5.1). A high EMT score indicates a mesenchymal-like phenotype, which is often associated with cancer metastasis. In contrast, a low EMT score corresponds to an epithelial-like phenotype, typically related to cell adhesion and a reduced likelihood of migration and invasion.

### CNV score

CNV scores were computed using the “tl.infercnv” function of infercnvpy package (Python package,v 0.4.2, https://github.com/icbi-lab/infercnvpy) . The core idea is to compare single-cell RNA sequencing data against reference normal cell populations to identify gene expression changes indicative of CNV. This method helps in detecting large-scale chromosomal alterations by assessing expression deviations.

### Tumor stemness

To calculate the tumor stemness score, we collected literature on stemness marker genes for various types of cancer and calculated the gene activity score to represent tumor stemness^28,34,45-48^ (**Supplementary Data 20**). Cells with higher scores are more likely to be occult metastatic cells.

### Metastatic enrichment score

To calculate metastatic enrichment score, we select the metastasis-related pathway in Gene Ontology (GO) term and calculated the gene activity score to represent tumor stemness. Cells with higher scores are more likely to be metastatic cells.

### Overview score

In each dataset used for computational evaluation, we rank five tools across EMT, CNV and tumor stemness indexes. A higher rank corresponds to a higher score. The highest possible score in a single dataset is 15, while the lowest is 3.

### True positive rate (TPR)

*TPR* refers to the proportion of actual positive cases (patients with metastasis) that are correctly identified by the model. It is defined as follows:

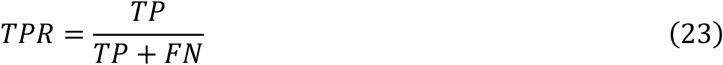

where:

- *TP* (True Positive) is the number of patients who have metastasis and are correctly classified as having metastasis.
- *FN* (False Negative) is the number of patients who have metastasis but are incorrectly classified as not having metastasis.

### False positive rate (FPR)

*FPR* refers to the proportion of actual negative cases (patients without metastasis) that are incorrectly classified as positive by the model. It is defined as follows:

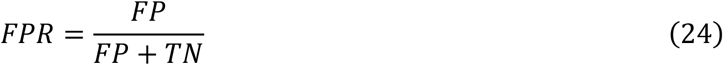

where:

- *FP* (False Positive) is the number of patients who do not have metastasis but are incorrectly classified as having metastasis.
- *TN* (True Negative) is the number of patients who do not have metastasis and are correctly classified as not having metastasis.

### Trajectory analysis

We used the “orderCells” function from the Monocle2^49^ (R package, v2.26.0) to infer cell developmental trajectories and rank the cells accordingly.

### Simulated TF knockout

We utilized the CellOracle^24^ (Python package, v0.12.0) to perform simulated transcription factor TF knockout analyses. First, we constructed gene regulatory networks using the “celloracle.data.load_human_promoter_base_GRN” function, which provides pre-built promoter base GRNs for human cells. Next, we applied the “celloracle.simulate_shift” function to simulate the knockout of specific TFs. This function recalculates the predicted gene expression profiles by removing the regulatory influence of the selected TFs, allowing us to assess the potential impact of the TF knockout on downstream gene expression and cellular behavior.

### Statistics and reproducibility

#### Statistics

All statistical analyses were performed in R version 4.2.1 (ggpubr 0.6.0, survival 3.8.3, and survminer 0.5.0) and Python version 3.8.0 (scipy 1.9.1). All boxplots display the median as the central line, the interquartile range (IQR; 25th to 75th percentile) as the box and outliers (points beyond 1.5 times the IQR) as dots outside the whiskers. Both line plots with error bands and bar plots with error bars display mean values, with the error bands and error bars representing the standard deviation above and below the means.

#### Experimental

The transwell migration assay was performed in three independent biological replicates, with each group containing six technical replicates per experiment. All experiments yielded consistent results, demonstrating reproducibility.

#### Model

To evaluate the stability and reproducibility of EmitGCL, we randomly selected a dataset from each cancer type and then repeated the algorithm 20 times using the default parameters. Finally, we calculated the ARI between the different runs for each selected dataset within each cancer type to assess the consistency of the clustering results.

### Clinical data integration

To integrate the data, we first collected the intersection of genes from five batches of uncorrected expression data and then filtered out genes do not present in the intersection across all datasets. We then integrated batch information from the different datasets into a single AnnData object. The following clinical data were added to the combined dataset: batch information (Batch: Chin, Caldas, Desmedt, Miller, TCGA), time information (Time), and label information (Label: high metastatic potential ≤ 5), low metastatic potential (Time > 5)).

### Clinical differential gene screening

Differentially expressed genes (DEGs) were identified within each batch using the Wilcoxon rank-sum test based on the label. For each batch, genes with p < 0.05 and logFC > 0 were selected as DEGs. The significant genes were then divided according to their batch, and an UpSet plot was generated to identify common DEGs across multiple batches. Common genes included *GATA3* (shared by Caldas and Chin) and *AZGP1* (shared by Caldas and Desmedt). Finally, *GATA3* and *AZGP1* were selected as the DEGs for benchmarking.

### Batch effect correction

Batch effects in the integrated data were corrected using the ComBat^50^ method, resulting in corrected expression data for further analysis.

### Model feature selection

The expression data of the corrected DEGs, *GATA3* and *AZGP1*, were extracted for model feature validation. Models were established using these feature data to analyze the relationship between genes in the training set. The model’s accuracy and predictive performance were evaluated across different gene combinations using random seeds for training. To visualize the results, we plotted the mean and maximum accuracy of each gene combination and created Kaplan-Meier survival curves to analyze the survival status of patients with early or late metastasis based on gene combinations. Finally, *HSP90AA1* and *HSP90AB1* were identified as the final model features.

### Classification model performance evaluation

The expression data of the corrected DEGs, *GATA3* and *AZGP1*, were used to evaluate the classification model’s performance. We focused on samples with late metastasis (Time > 5) for *GATA3* and *AZGP1* expression, using various random seeds to assess classification accuracy. The scatter plot of *GATA3* and *AZGP1* gene expression was visualized, and the corresponding classification boundaries were drawn. The classification performance was illustrated based on the identified features for samples in early and late metastasis stages.

### Clustering model performance evaluation

For clustering model performance evaluation, we extracted the expression data of the corrected DEGs, *GATA3* and *AZGP1*. K-means clustering was employed to group the data into two clusters. The clustering accuracy was calculated, and the results were visualized. The clustering results were displayed, and the distribution of clusters based on gene expression was analyzed.

## Data availability

Previously published scRNA-seq datasets were used for benchmarking and case studies in this study. Breast cancer scRNA-seq datasets were obtained from GEO under accession numbers GSE167036 and GSE180286. Lung cancer datasets were obtained from GEO under accession number GSE198099. Head and neck cancer datasets were obtained from GEO under accession number GSE188737. Pancreatic cancer datasets were obtained from GEO under accession number GSE197177. Papillary thyroid cancer datasets were obtained from GEO under accession number GSE214148. Nasopharyngeal carcinoma scRNA-seq datasets were obtained from the GSA for Human database under accession number HRA000036. Additional previously published RNA-seq datasets from UCSC were used for breast cancer studies, including datasets from Chin (2006), Miller (2005), Desmedt (2007), Wang (2005), and Caldas (2007). All datasets were reprocessed and analyzed as described in the Methods section. Details of data information can be found in **Supplementary Data 1**. Source data are provided in this paper.

## Code availability

The code supporting this study is not publicly available. However, the source code is provided as a Code.zip file in the supplementary materials for reviewer access.

## Acknowledgments

This work was supported by research grants P01CA278732 (Z.L. and Q.M.) from the National Institutes of Health. This work was supported by the Pelotonia Institute of Immuno-Oncology (PIIO). The content is solely the responsibility of the authors and does not necessarily represent the official views of the PIIO.

## Author contributions

Q.M. conceived the basic idea. X.W. designed the algorithm, conducted the case study, and wrote the manuscript. M.D. carried out benchmark experiments and wrapped the code. P.S. performed survival analysis. J.L., J.J., and X.G. designed and conducted cell line study. J.K. provided biological insights for this study. P.S., K.H., Z.L., and D.P.C. provided clinical rationale for this study. D.X. provided deep learning knowledge. B.L., H.C., Y.S., and W.W. contributed to methodological guidance and discussion interpretation. G.W., C.Z., S.C. contributed expert insights during manuscript preparation and provided critical feedback

## References

1 Fares, J., Fares, M. Y., Khachfe, H. H., Salhab, H. A. & Fares, Y. Molecular principles of metastasis: a hallmark of cancer revisited. Signal Transduct Target Ther 5, 28 (2020). 10.1038/s41392-020-0134-x

2 Lusby, R., Dunne, P. & Tiwari, V. K. Tumour invasion and dissemination. Biochem Soc Trans 50, 1245–1257 (2022). 10.1042/BST20220452

3 Dieci, M. V. et al. Metastatic site patterns by intrinsic subtype and HER2DX in early HER2-positive breast cancer. J Natl Cancer Inst 116, 69–80 (2024). 10.1093/jnci/djad179

4 Zhan, Q. et al. New insights into the correlations between circulating tumor cells and target organ metastasis. Signal Transduct Target Ther 8, 465 (2023). 10.1038/s41392-023-01725-9

5 Luo, L. et al. Single-cell RNA sequencing identifies molecular biomarkers predicting late progression to CDK4/6 inhibition in patients with HR+/HER2-metastatic breast cancer. Mol Cancer 24, 48 (2025). 10.1186/s12943-025-02226-9

6 Zhou, J. et al. PLUS: Predicting cancer metastasis potential based on positive and unlabeled learning. PLoS Comput Biol 18, e1009956 (2022). 10.1371/journal.pcbi.1009956

7 Wysong, A. et al. Validation of a 40-gene expression profile test to predict metastatic risk in localized high-risk cutaneous squamous cell carcinoma. J Am Acad Dermatol 84, 361–369 (2021). 10.1016/j.jaad.2020.04.088

8 Barca-Hernando, M. et al. Occult cancer in patients with unprovoked venous thromboembolism: Rationale, design, and methods of the ValRIETEs study and the SOME-RIETE trial. Am Heart J (2025). 10.1016/j.ahj.2025.02.004

9 Ng, L., Poon, R. T. & Pang, R. Biomarkers for predicting future metastasis of human gastrointestinal tumors. Cell Mol Life Sci 70, 3631–3656 (2013). 10.1007/s00018-013-1266-8

10 Steeg, P. S. Targeting metastasis. Nat Rev Cancer 16, 201–218 (2016). 10.1038/nrc.2016.25

11 Jaeger, J. et al. Gene expression signatures for tumor progression, tumor subtype, and tumor thickness in laser-microdissected melanoma tissues. Clin Cancer Res 13, 806–815 (2007). 10.1158/1078-0432.CCR-06-1820

12 Sun, S., Shi, R., Xu, L. & Sun, F. Identification of heterogeneity and prognostic key genes associated with uveal melanoma using single-cell RNA-sequencing technology. Melanoma Res 32, 18–26 (2022). 10.1097/CMR.0000000000000783

13 Smith, A. P., Hoek, K. & Becker, D. Whole-genome expression profiling of the melanoma progression pathway reveals marked molecular differences between nevi/melanoma in situ and advanced-stage melanomas. Cancer Biol Ther 4, 1018–1029 (2005). 10.4161/cbt.4.9.2165

14 Winnepenninckx, V. et al. Gene expression profiling of primary cutaneous melanoma and clinical outcome. J Natl Cancer Inst 98, 472–482 (2006). 10.1093/jnci/djj103

15 Forouzandeh, A., Rutar, A., Kalmady, S. V. & Greiner, R. Analyzing biomarker discovery: Estimating the reproducibility of biomarker sets. PLoS One 17, e0252697 (2022). 10.1371/journal.pone.0252697

16 Safari, F., Kehelpannala, C., Safarchi, A., Batarseh, A. M. & Vafaee, F. Biomarker Reproducibility Challenge: A Review of Non-Nucleotide Biomarker Discovery Protocols from Body Fluids in Breast Cancer Diagnosis. Cancers (Basel) 15 (2023). 10.3390/cancers15102780

17 Jiang, L., Chen, H., Pinello, L. & Yuan, G. C. GiniClust: detecting rare cell types from single-cell gene expression data with Gini index. Genome Biol 17, 144 (2016). 10.1186/s13059-016-1010-4

18 Ren, L. et al. Single cell RNA sequencing for breast cancer: present and future. Cell Death Discov 7, 104 (2021). 10.1038/s41420-021-00485-1

19 Ma, A. et al. Single-cell biological network inference using a heterogeneous graph transformer. Nat Commun 14, 964 (2023). 10.1038/s41467-023-36559-0

20 Gui, Y., He, X., Yu, J. & Jing, J. Artificial Intelligence-Assisted Transcriptomic Analysis to Advance Cancer Immunotherapy. J Clin Med 12 (2023). 10.3390/jcm12041279

21 Ilangovan, H. et al. Harmonizing heterogeneous transcriptomics datasets for machine learning-based analysis to identify spaceflown murine liver-specific changes. NPJ Microgravity 10, 61 (2024). 10.1038/s41526-024-00379-3

22 Ma, Q. & Xu, D. Deep learning shapes single-cell data analysis. Nat Rev Mol Cell Biol 23, 303–304 (2022). 10.1038/s41580-022-00466-x

23 Wang, S. et al. UCSCXenaShiny: an R/CRAN package for interactive analysis of UCSC Xena data. Bioinformatics 38, 527–529 (2022). 10.1093/bioinformatics/btab561

24 Kamimoto, K. et al. Dissecting cell identity via network inference and in silico gene perturbation. Nature 614, 742–751 (2023). 10.1038/s41586-022-05688-9

25 Wang, X. et al. MarsGT: Multi-omics analysis for rare population inference using single-cell graph transformer. Nat Commun 15, 338 (2024). 10.1038/s41467-023-44570-8

26 Stuart, T. et al. Comprehensive Integration of Single-Cell Data. Cell 177, 1888-1902.e1821 (2019). 10.1016/j.cell.2019.05.031

27 Wegmann, R. et al. CellSIUS provides sensitive and specific detection of rare cell populations from complex single-cell RNA-seq data. Genome Biol 20, 142 (2019). 10.1186/s13059-019-1739-7

28 Quah, H. S. et al. Single cell analysis in head and neck cancer reveals potential immune evasion mechanisms during early metastasis. Nat Commun 14, 1680 (2023). 10.1038/s41467-023-37379-y

29 Liu, Y. M. et al. Combined Single-Cell and Spatial Transcriptomics Reveal the Metabolic Evolvement of Breast Cancer during Early Dissemination. Adv Sci (Weinh) 10, e2205395 (2023). 10.1002/advs.202205395

30 Zhou, W. et al. Cancer Stemness Online: A Resource for Investigating Cancer Stemness and Associations with Immune Response. Genomics Proteomics Bioinformatics 22 (2024). 10.1093/gpbjnl/qzae058

31 Thomassen, M., Tan, Q. & Kruse, T. A. Gene expression meta-analysis identifies metastatic pathways and transcription factors in breast cancer. BMC Cancer 8, 394 (2008). 10.1186/1471-2407-8-394

32 Prizment, A. E. et al. Circulating Beta-2 Microglobulin and Risk of Cancer: The Atherosclerosis Risk in Communities Study (ARIC). Cancer Epidemiol Biomarkers Prev 25, 657–664 (2016). 10.1158/1055-9965.EPI-15-0849

33 Gutschner, T. et al. The noncoding RNA MALAT1 is a critical regulator of the metastasis phenotype of lung cancer cells. Cancer Res 73, 1180–1189 (2013). 10.1158/0008-5472.CAN-12-2850

34 Xu, K. et al. Single-cell RNA sequencing reveals cell heterogeneity and transcriptome profile of breast cancer lymph node metastasis. Oncogenesis 10, 66 (2021). 10.1038/s41389-021-00355-6

35 Harvey, K. F., Zhang, X. & Thomas, D. M. The Hippo pathway and human cancer. Nat Rev Cancer 13, 246–257 (2013). 10.1038/nrc3458

36 Fu, M. et al. The Hippo signalling pathway and its implications in human health and diseases. Signal Transduct Target Ther 7, 376 (2022). 10.1038/s41392-022-01191-9

37 Zheng, Y. & Pan, D. The Hippo Signaling Pathway in Development and Disease. Dev Cell 50, 264–282 (2019). 10.1016/j.devcel.2019.06.003

38 Dugina, V. et al. Imbalance between Actin Isoforms Contributes to Tumour Progression in Taxol-Resistant Triple-Negative Breast Cancer Cells. Int J Mol Sci 25 (2024). 10.3390/ijms25084530

39 Keenan, A. B. et al. ChEA3: transcription factor enrichment analysis by orthogonal omics integration. Nucleic Acids Res 47, W212–W224 (2019). 10.1093/nar/gkz446

40 Ren, F. J., Cai, X. Y., Yao, Y. & Fang, G. Y. JunB: a paradigm for Jun family in immune response and cancer. Front Cell Infect Microbiol 13, 1222265 (2023). 10.3389/fcimb.2023.1222265

41 Casalino, L., Talotta, F., Matino, I. & Verde, P. FRA-1 as a Regulator of EMT and Metastasis in Breast Cancer. Int J Mol Sci 24 (2023). 10.3390/ijms24098307

42 Chan, H. L. et al. Polycomb complexes associate with enhancers and promote oncogenic transcriptional programs in cancer through multiple mechanisms. Nat Commun 9, 3377 (2018). 10.1038/s41467-018-05728-x

43 Zhang, L. et al. Z β β -FAK activation. Nat Commun 13, 2543 (2022). 10.1038/s41467-022-30105-0

44 Andreatta, M. & Carmona, S. J. UCell: Robust and scalable single-cell gene signature scoring. Comput Struct Biotechnol J 19, 3796–3798 (2021). 10.1016/j.csbj.2021.06.043

45 Liu, T. et al. Single cell profiling of primary and paired metastatic lymph node tumors in breast cancer patients. Nat Commun 13, 6823 (2022). 10.1038/s41467-022-34581-2

46 Zhang, S. et al. Single cell transcriptomic analyses implicate an immunosuppressive tumor microenvironment in pancreatic cancer liver metastasis. Nat Commun 14, 5123 (2023). 10.1038/s41467-023-40727-7

47 Chen, W. et al. Single-cell RNA-seq reveals MIF-(CD74 + CXCR4) dependent inhibition of macrophages in metastatic papillary thyroid carcinoma. Oral Oncol 148, 106654 (2024). 10.1016/j.oraloncology.2023.106654

48 Xu, D. et al. Single-cell sequencing analysis reveals the dynamic tumour ecosystems of primary and metastatic lymph nodes in nasopharyngeal carcinoma. J Cell Mol Med 28, e70137 (2024). 10.1111/jcmm.70137

49 Qiu, X. et al. Reversed graph embedding resolves complex single-cell trajectories. Nat Methods 14, 979–982 (2017). 10.1038/nmeth.4402

50 Johnson, W. E., Li, C. & Rabinovic, A. Adjusting batch effects in microarray expression data using empirical Bayes methods. Biostatistics 8, 118–127 (2007). 10.1093/biostatistics/kxj037

